# AT-752, a double prodrug of a guanosine nucleotide analog, inhibits yellow fever virus in a hamster model

**DOI:** 10.1101/2021.10.25.465665

**Authors:** Kai Lin, Steven S. Good, Justin G. Julander, Abbie Weight, Adel Moussa, Jean-Pierre Sommadossi

**Affiliations:** Atea Pharmaceuticals, Inc., Boston, Massachusetts, United States of America; Institute for Antiviral Research, Utah State University, Logan, Utah, USA

**Author notes:** Corresponding author (KL).

**Keywords:** AT-752, yellow fever virus, flavivirus, hamster, antiviral

## Abstract

Yellow fever virus (YFV) is a zoonotic pathogen re-emerging in parts of the world, causing a viral hemorrhagic fever associated with high mortality rates. While an effective vaccine is available, having an effective antiviral against YFV is critical against unexpected outbreaks, or when vaccination is not recommended. We have previously identified AT-281, the free base of AT-752, an orally available double prodrug of a guanosine nucleotide analog, as a potent inhibitor of YFV *in vitro*, with a 50% effective concentration (EC_50_) of 0.31 μM. In hamsters infected with YFV (Jimenez strain), viremia rose about 4 log_10_-fold and serum alanine aminotransferase (ALT) 2-fold compared to sham-infected animals. Treatment with 1000 mg/kg AT-752 for 7 days, initiated 4 h prior to viral challenge, reduced viremia to below the limit of detection by day 4 post infection (pi) and returned ALT to normal levels by day 6 pi. When treatment with AT-752 was initiated 2 days pi, the virus titer and ALT dropped >2 log_10_ and 53% by day 4 and 6 pi, respectively. In addition, at 21 days pi, 70 – 100% of the infected animals in the treatment groups survived compared to 0% of the untreated group (p<0.001). Moreover, *in vivo* formation of the active triphosphate metabolite AT-9010 was measured in the animal tissues, with the highest concentrations in liver and kidney, organs that are vulnerable to the virus. The demonstrated *in vivo* activity of AT-752 suggests that it is a promising compound for clinical development in the treatment of YFV infection.

**Author summary:** Yellow fever virus (YFV) is transmitted by mosquitoes, and its infection can lead to a lethal viral hemorrhagic fever associated with liver damage. While an effective vaccine is available, in places where the vaccination rate is low, in the event of an unexpected outbreak, or where vaccination is not recommended individually, having an effective antiviral treatment is critical. We previously reported that the nucleotide analog prodrug AT-752 potently inhibited the YFV in cultured cells. Here we showed that in hamsters infected with YFV, oral treatment with 1000 mg/kg AT-752 for 7 days reduced the production of infectious virus particles in the blood, and decreased serum alanine aminotransferase, a marker of liver damage, to levels measured in uninfected animals. In addition, at 21 days after infection, 70 – 100% of the infected animals in the treatment groups survived compared to 0% in the untreated group. Moreover, the amount of the active metabolite formed from AT-752 was highest in the livers and kidneys of the treated animals, organs that are targeted by the virus. These results suggest that AT-752 is a promising compound to develop for the treatment of YFV infection.

## Introduction

Yellow fever virus (YFV) is one of the single-stranded, positive-sense RNA viruses of the Flaviviridae family. According to the World Health Organization (WHO), yellow fever is endemic in forty-seven countries in Africa and Central and South America, and half of the patients that develop severe symptoms from this mosquito-borne virus die within 7 – 10 days [1]. Infection from YFV is difficult to diagnose, but in severe cases, it affects the kidneys and liver, causing jaundice with the latter, hence the name “yellow fever”. Although there is an effective vaccine, it is not recommended for pregnant and lactating women, immune-compromised individuals nor those older than 60 years [2, 3], so there is a real need for an effective antiviral treatment. Moreover, this deadly virus is a threat in regions where the vaccine is under-utilized, resulting in unanticipated cases from outbreaks or emergence in areas that have previously not been affected by the virus, as observed by the recent large outbreaks and YFV emergence in Brazil and several African countries [4, 5] that have caused significant morbidity and mortality.

Currently, there are no approved antiviral therapies for the treatment of yellow fever. Drugs such as interferon [6] and ivermectin [7] are no longer being pursued, however, sofosbuvir [2, 8] is being examined for the purpose of treating this malady until better options become available. There have been many attempts by other investigators to develop direct-acting antiviral (DAA) medications against yellow fever, from a benzodiazepine compound that targets the nonstructural protein (NS) 4B which is thought to anchor the viral replication complex [9] to nucleoside analogs that were effective in animal models [10, 11], but no compound has successfully demonstrated efficacy in human testing. Nucleoside analogs are promising potential drugs for DAA development because they target the highly conserved NS5 protein and have shown potent antiviral activity against YFV *in vitro* [3]. Flavivirus NS5 proteins have two functional domains, an N-terminal methyltransferase (MTase) domain and a C-terminal RNA-dependent RNA polymerase (RdRp) domain, which play essential roles in viral RNA replication, transcription and capping, so disrupting the activity of YFV NS5 would inhibit viral replication [4].

We have developed a 2’-fluoro-2’-C-methyl guanosine nucleotide prodrug AT-752, and we have reported that its free base AT-281 has potent *in vitro* antiviral activity against YFV and several other flaviviruses [12]. Against YFV, the effective concentration of AT-281, the free base of the salt form AT-752, required to achieve 50% inhibition (EC_50_) of the virus-induced cytopathic effect (CPE) was 0.31 μM, with an EC_90_ of 0.26 μM in reducing viral yield. In addition, this nucleotide analog functions by having the active intracellular triphosphate metabolite AT-9010 (see putative pathway, Fig 1) inhibit NS5, causing termination of viral RNA synthesis [12]. We now show the efficacy of AT-752 against YFV in a hamster model.

**Fig 1.**
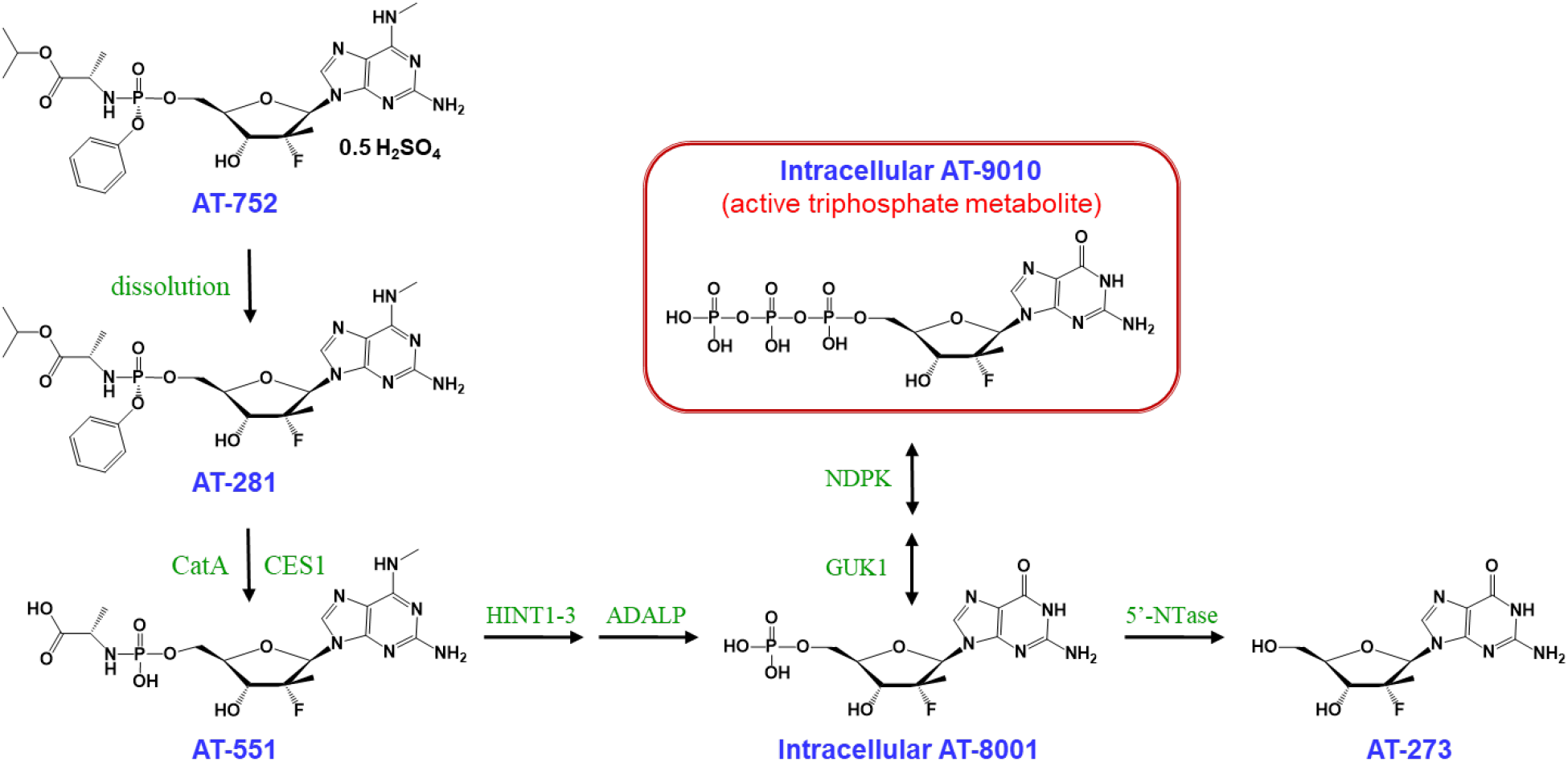
AT-752 and its putative metabolic pathway to the pharmacologically active metabolite AT-9010. The metabolite AT-273 is a plasma surrogate for AT-9010.

## Methods

### Cells, viruses and test compounds

Vero 76 cells (American Type Culture Collection, Manassas, VA) used to verify viral titers were maintained in Dulbecco’s Modified Eagle medium (DMEM) supplemented with 10% fetal bovine serum (FBS) and 100 μg/mL penicillin and 100 μg/mL streptomycin (Lonza, Walkersville, MD) at 37°C in an atmosphere of 5% CO_2_ and ≥95% humidity. The yellow fever virus (YFV Jimenez hamster-adapted strain) was obtained as a generous gift from R. B. Tesh (University of Texas Medical Branch, Galveston, TX) and stock was prepared by a further passage in hamsters (USU V#2653). AT-752, its free base AT-281, and metabolites AT-551 and AT-273 were prepared for Atea Pharmaceuticals by Topharman Shanghai Co., Ltd., Shanghai, China. AT-9010 and the triphosphate internal standards used to quantify AT-9010 were synthesized by NuBlocks (Oceanside, CA). Stock solutions were prepared in DMSO and stored at −20°C.

### Animal welfare

The two animal studies described herein were conducted in AAALAC-accredited Laboratory Animal Research Centers which strictly comply with the USDA Animal Welfare Act and follow the PHS and NIH Policy of Humane Care and Use of Laboratory Animals, and the Guide for the Care and Use of Laboratory Animals, National Research Council – ILAR, Revised 2011. The study where hamsters were infected with YFV was conducted at Utah State University (USU, Logan, UT) in full compliance with protocols that were reviewed and approved by USU Institutional Animal Care and Use Committee (IACUC) prior to study initiation (IACUC number #10010), and hamsters were assessed and monitored throughout the study by members of the USU veterinary staff in accordance with AAALAC. The pharmacokinetic study was conducted in full compliance with the approved protocol at WuXi AppTec (Shanghai, China).

### Pharmacokinetics (PK) and tissue distribution of AT-752 in hamsters

Fifteen male Syrian golden hamsters (Beijing VITAL RIVER Laboratory Animals Co. Ltd., China) were set in 5 groups of 3 hamsters each and administered a dose of AT-752 at 500 mg/kg in 40% PEG400, 10% solutol HS15, 50% 100 mM citrate buffer, pH 4.5 (v/v) by oral gavage. Blood samples were collected from animals at 2 (group 1), 4 (group 2), 8 (group 3) and 12 h (group 4) post-dose. Serial blood samples were collected from animals in Group 5 at 0.5, 1, 2, 4, 8, 12 and 24 h post dose. For all samples, plasma was separated in EDTA and 5 μL dichlorvos (2 mg/mL; stabilizing agent to prevent *in vitro* hydrolysis of the ester moiety of AT-281 by blood esterases) and stored at −60°C. Concentrations of AT-281, and metabolites AT-551 and AT-273 were determined in the plasma by LC/MS/MS.

After the blood collection at each terminal timepoint, the animals were anesthetized via isoflurane. Liver, kidney, lung and brain were removed immediately, wrapped quickly with aluminum foil, and dropped into liquid nitrogen within 20 seconds. Once frozen, tissue samples were placed in a pre-labeled bag, and stored at −60°C until further processing, at which point, they were weighed and homogenized using a Bead beater in 5 volumes ice-cold homogenization buffer containing 70% methanol, 30% 268 mM K_2_EDTA (final pH 7.8), and internal standards. All homogenates were stored at −80°C until analyzed for concentrations of AT-281, and metabolites, AT-551, AT-273 and AT-9010 using LC/MS/MS.

#### LC-MS/MS analysis of AT-281, AT-551, AT-273 and AT-9010

Plasma samples were prepared for MS analysis by adding internal standards and extracting with 20 volumes chilled MeOH/ACN (75:25, v/v). After vortex mixing (800 rpm for 10 min) and centrifugation (3220 *g*, 15 min, 4°C), the supernatants (25 μL) were diluted with an equal volume of H_2_O, mixed and spun again. For the tissue homogenates, after adding internal standards, 40 μL was mixed by vortex with 400 μL MeOH/ACN (75:25, v/v), and centrifuged (3220 *g*, 15 min, 4°C). Aliquots of the supernatants (25 μL) were diluted with an equal volume of H_2_O, mixed and spun again. To measure AT-281 and plasma metabolites AT-551 and AT-273, 4 μL samples were injected onto an Acquity Gemini C18 (50 x 4.6 mm), 5 μm UPLC column with a Sciex Triple Quad 6500 mass spectrometer (ESI positive ion, MRM mode). A binary nonlinear gradient with mobile phases A (0.1% formic acid in water) and B (0.1% formic acid in ACN) were used to elute samples at 0.8 mL/min, with a run time of 5 min. For AT-9010, 40 μL of 20 mM NH_4_OAC buffer (pH 8.4) was added to the homogenate (40 μL) along with internal standards, and samples were lysed with 160 μL MeOH and 40 mM dibutylammonium acetate (DBAA), mixed by vortex and centrifuged (3220 *g*, 15 min, 4°C). Aliquots of the supernatants (35 μL) were dried under nitrogen and reconstituted in H_2_O, then mixed and spun again before injecting 15 μL onto an Acquity BEH C18 (50 x 2.1 mm), 1.7 μm UPLC column with an API 14000 mass spectrometer (ESI negative ion, MRM mode). A binary nonlinear gradient with mobile phases A (0.001% NH_3_·H_2_O, 0.18 mM DBAA in H_2_O) and B (10 mM N, N-dimethyl-hexylamine, 3 mM NH_4_OAc in ACN/H_2_O (50:50, v/v)) were used to elute samples at 0.5 mL/min, with a run time of 3 min. Standards in 50% MeOH were used for calibration. Ions monitored were m/z 538.2/158.8 (AT-9010), 582.3/330.1 (AT-281), 464.2/165.1 (AT-551) and 300.1/152.1 (AT-273). Internal standards (ISS) as described previously [13] were used to correct for variations in recovery.

#### PK Data Analysis

Plasma and tissue concentrations of AT-281, AT-551 and AT-273 were subjected to non-compartmental pharmacokinetic analysis using Phoenix WinNonlin software (version 6.3, Pharsight, Mountain View, CA). The linear/log trapezoidal rule was applied in obtaining the PK parameters.

### AT-752 treatment against YFV in a hamster model

Eighty female Syrian golden hamsters (LVG/Lak strain, Charles River), 90 – 110 g body weight, were divided into nine groups as shown in Table 1. On Day 0, all the treatment animals (Groups 1 – 6) were inoculated with YFV, 200 CCID_50_ per hamster in 0.1 mL volume via bilateral intraperitoneal injection. This dose was approximately 6x the LD_50_ in hamsters. Group 1 – 4 animals received vehicle [(PEG400 (40%, v/v)/ Solutol HS15 (10%, v/v)/100 mM Citrate buffer pH 4.5 (50%, v/v)], 1000, 300 and 100 mg/kg of AT-752 respectively, by oral gavage twice daily (BID) for 7 consecutive days starting 4 h prior to challenge. Group 5 received an oral dose of 1000 mg/kg AT-752 two days post challenge (pi), BID for 7 consecutive days while Group 6 were administered 50 mg/kg ribavirin (RIBA), 4 h prior to challenge by intraperitoneal injection, BID for 7 consecutive days. Two control groups were sham infected 4 h after being given 1000 mg/kg AT-752 or vehicle orally, followed by BID dosing for 7 consecutive days. The final control group did not receive virus or treatment. The hamsters were monitored for 21 and 18 days post virus challenge for survival and weight change, respectively. Blood samples were taken ante mortem via ocular sinus bleed on Day 4 pi for quantification of virus by infectious cell culture assay to determine the 50% cell culture infectious dose (CCID_50_) and on Day 6 pi to measure alanine aminotransferase (ALT).

**Table 1.** Treatment groups for the hamster study.

| Test compound | n/group | Dose (mg/kg/d) | Treatment Schedule | Virus |
| --- | --- | --- | --- | --- |
| AT-752 | 10 | 1,000 | 0.5 mL, p.o., bid X 7 d beg. -4 h | YFV |
| AT-752 | 10 | 300 | 0.5 mL, p.o., bid X 7 d beg. -4 h | YFV |
| AT-752 | 10 | 100 | 0.5 mL, p.o., bid X 7 d beg. -4 h | YFV |
| AT-752 | 10 | 1,000 | 0.5 mL, p.o., bid X 7 d beg. 2d pi | YFV |
| Ribavirin | 10 | 50 | 0.5 mL, i.p., bid X 7 d beg. -4 h | YFV |
| Vehicle | 15 | -- | 0.5 mL, p.o. bid X 7 d beg. -4 h | YFV |
| AT-752 | 5 | 1,000 | 0.5 mL, p.o., bid X 7 d beg. -4h | Sham |
| Vehicle | 5 | -- | 0.5 mL, p.o., bid X 7 d beg. -4 h | Sham |
| Control | 5 | -- | -- | NA |

Syrian golden hamsters were administered AT-752 (100 mg/mL) or vehicle orally, or ribavirin (RIBA) by intraperitoneal injection, BID for 7 d, beginning 4 h prior or 2 d post infection with YFV (200 CCID<sub>50</sub> per animal). Three control groups (sham infected or no treatment) were included in the study.

#### Infectious cell culture assay

Virus titer was quantified using an infectious cell culture assay where a specific volume of serum was added to the first tube of a series of dilution tubes. Serial dilutions were made and added to Vero cells. Ten days later cytopathic effect (CPE) was used to identify the end-point of infection. Four replicates were used to calculate the 50% cell culture infectious doses (CCID_50_) per mL of plasma [10].

#### Serum alanine aminotransferase assay

Blood samples taken via ocular sinus bleed on Day 6 pi were centrifuged and serum collected. Alanine aminotransferase (ALT) was then measured using the ALT reagent from Teco Diagnostics (Anaheim, CA) in a protocol modified for 96-well plates. Briefly, 50 μl aminotransferase substrate was placed in each well of a 96-well plate, and 15 μl of sample was added at timed intervals. The samples were incubated at 37°C, after which 50 μl color reagent was added to each sample and incubated for 10 min as above. A volume of 200 μl of color developer was next added to each well and incubated for 5 min. The plate was then read on a spectrophotometer, and ALT concentrations were determined per manufacturer’s instructions.

#### Statistical analysis

Survival data were analyzed using the Wilcoxon log-rank survival analysis and all other statistical analyses were done using one-way ANOVA and a Dunnett multiple comparison (Prism 5, GraphPad Software, Inc).

## Results

### AT-752 has favorable pharmacokinetics in hamsters

The plasma pharmacokinetics (PK) and tissue distribution of AT-281 and its metabolites AT-551, the intermediate prodrug, and AT-273, the plasma surrogate for intracellular levels of the active triphosphate metabolite AT-9010, were determined in male Syrian golden hamsters after a single oral dose of AT-752 at 500 mg/kg (Table 2, Fig 2 and 3). In hamsters as in other rodents tested, the parent prodrug AT-281 was quickly absorbed and metabolized with the rapid appearance of its intermediate metabolite AT-551 in plasma, C_max_ 52.4 ± 23.5 nmol/mL at 0.5 h (Table 2). The prodrug was converted to AT-551 and AT-273, with plasma PK comparable to previously reported results of its congener AT-511 [13]. The plasma concentrations at 12 h post dose (or C_trough_, given that the dosing for the efficacy study was twice a day (BID) were 0.006 ± 0.002 nmol/mL for AT-281, 3.4 ± 1.7 nmol/mL for AT-551, and 0.6 ± 0.1 nmol/mL for AT-273. Concentrations of these three metabolites were also measured in the tissues collected – brain, lung, liver and kidney – up to 24 h post dose (Fig 2 and 3), although no AT-281 was detected in the lung and brain.

**Table 2.**
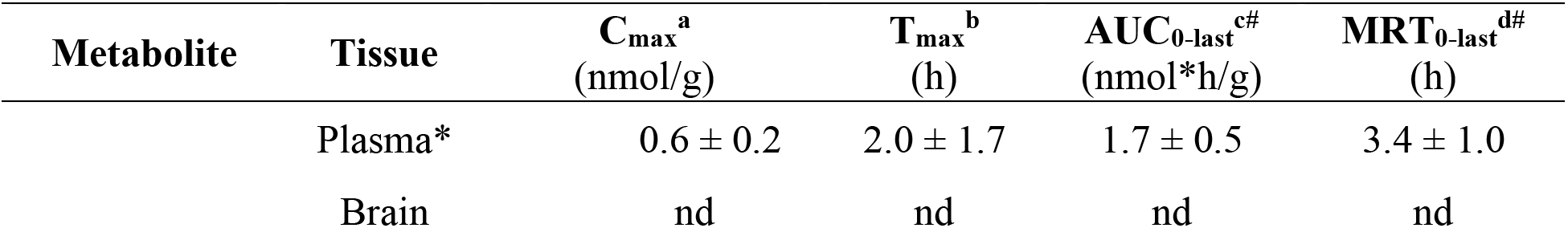
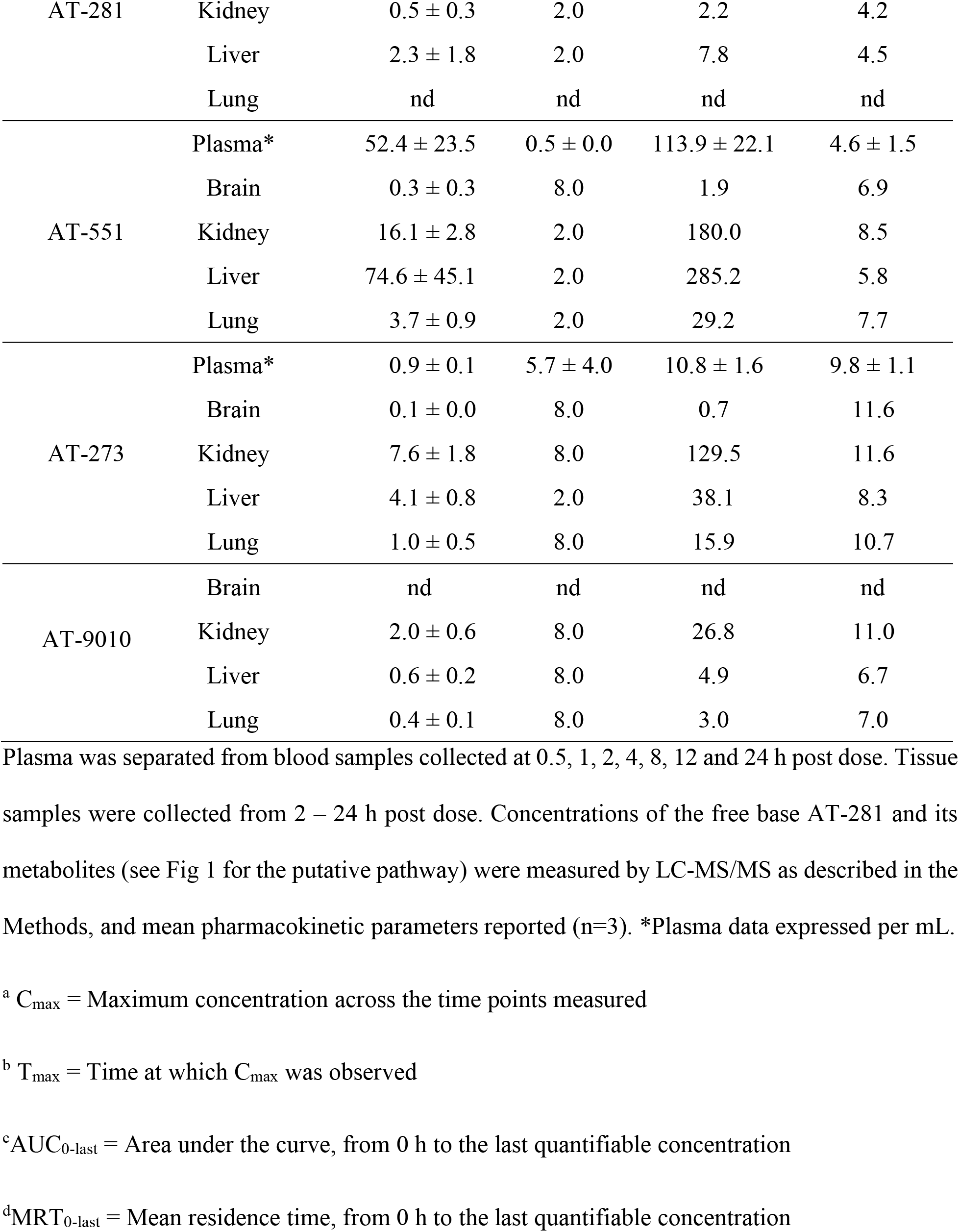
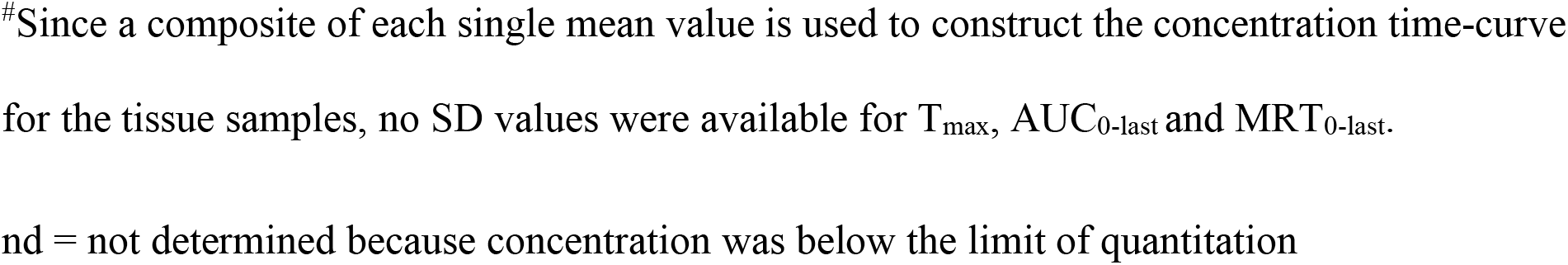
Pharmacokinetic parameters in male Syrian golden hamsters following oral administration of 500 mg/kg AT-752.

**Fig 2.**
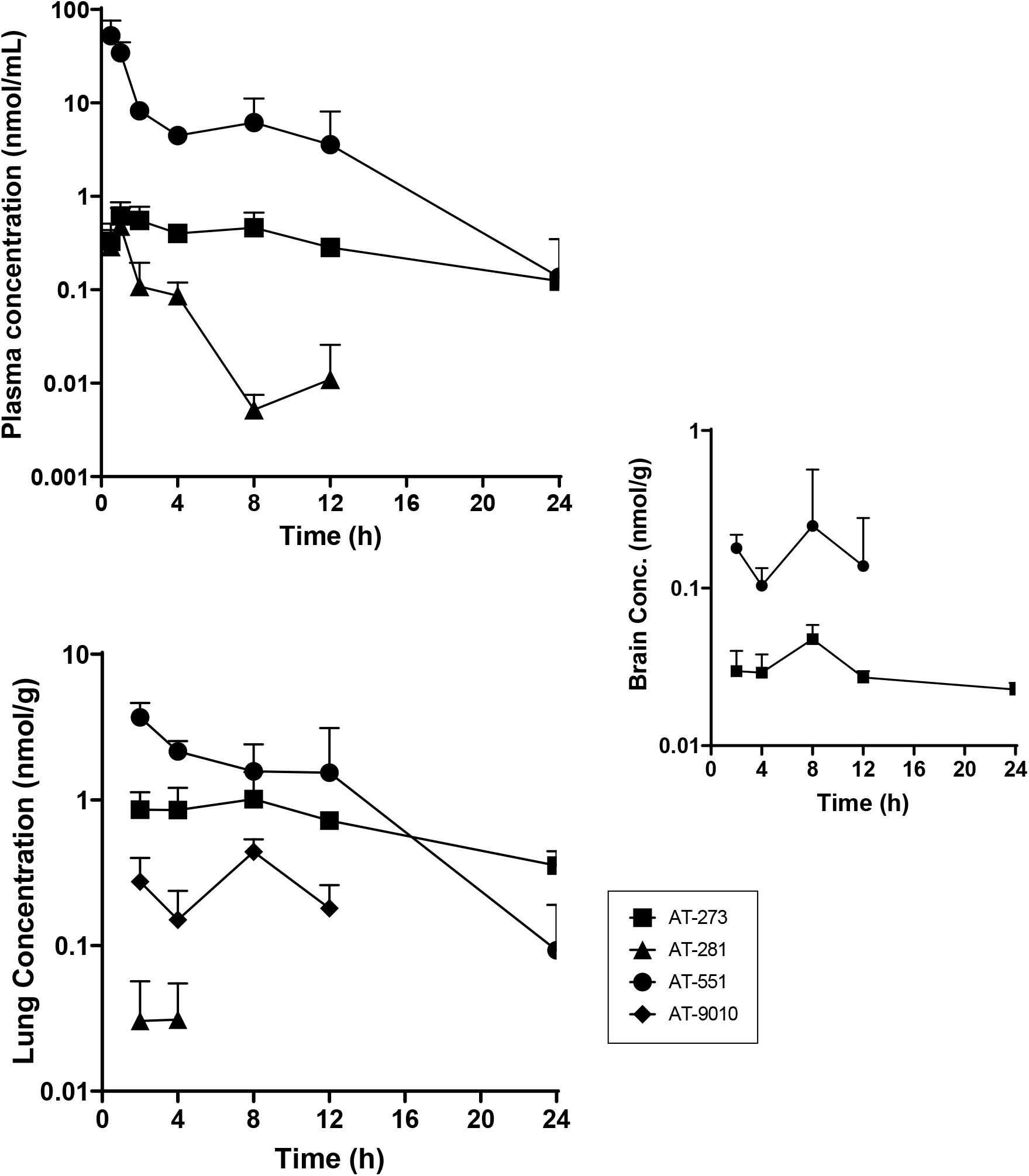
Profile of AT-281 and its metabolites in plasma, lung and brain tissue after a single oral dose of AT-752. Syrian golden male hamsters were administered 500 mg/kg AT-752. Samples were collected up to 24 h post dose and analyzed for AT-281 and its metabolites by LC-MS/MS. Data are expressed as mean ± SD (n= 3 per time point).

**Fig 3.**
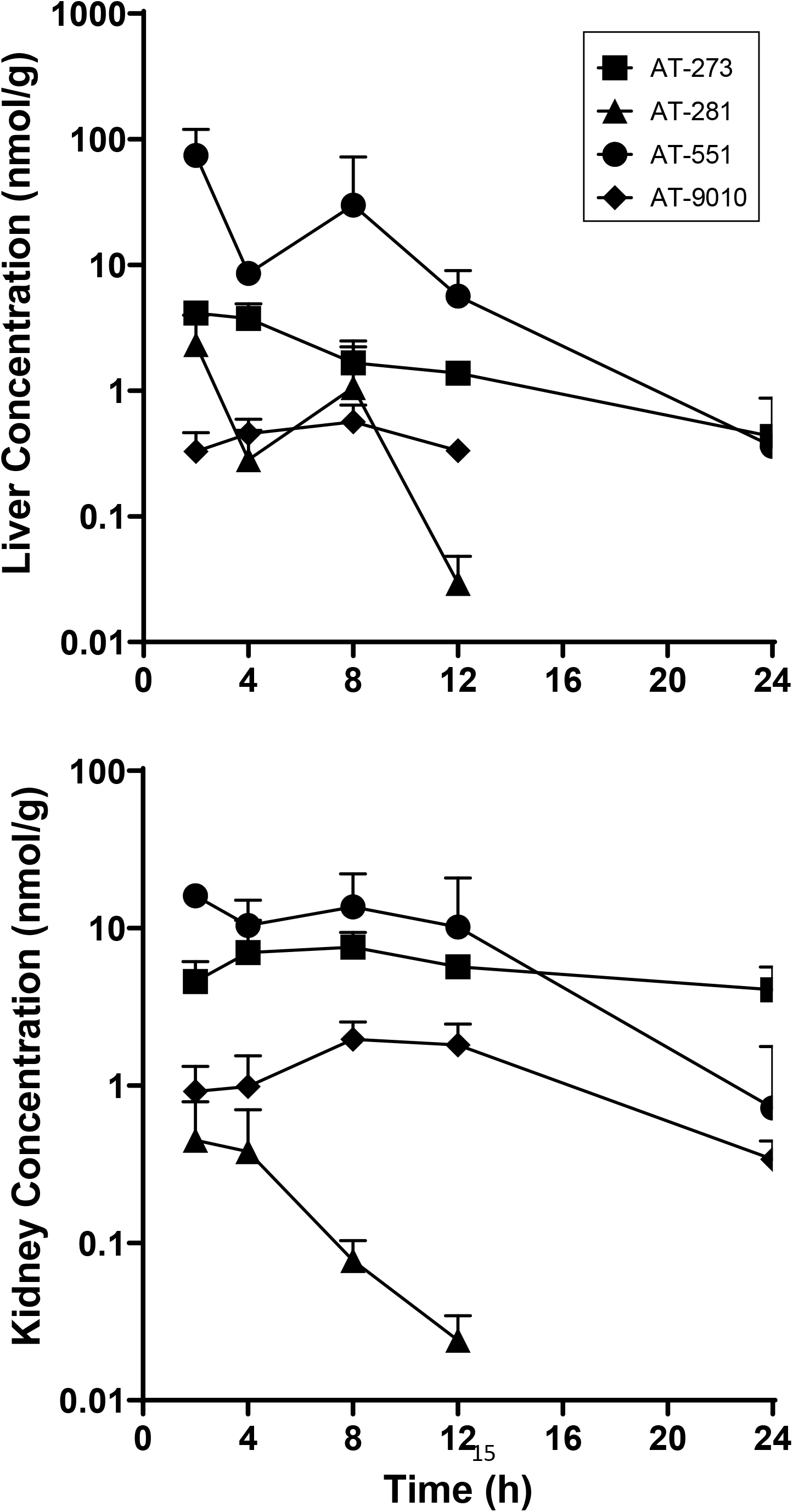
Profile of AT-281 and its metabolites in liver and kidney tissue after a single oral dose of AT-752. Syrian golden male hamsters were administered 500 mg/kg AT-752. Tissue samples were collected up to 24 h post dose and analyzed for AT-281 and its metabolites by LC-MS/MS. Data are expressed as mean ± SD (n= 3 per time point).

Liver and kidney tissue, which are most vulnerable in YFV infection, had the highest concentrations of AT-281 and its metabolites. In addition, concentrations of the active triphosphate AT-9010 were measured in lung, liver and kidney tissues, indicating *in vivo* formation of the intracellular metabolite (Fig 2 and 3), with mean residence times (MRT_0-last_) between 7 and 11 h (Table 2). There was no AT-9010 detected in the brain samples but given the low levels of the other metabolites measured in that tissue (Fig 2), it is likely that the prodrug does not easily cross the blood-brain barrier.

### AT-752 reduces viremia and improves disease outcomes of YFV-infected hamsters

To test the efficacy of AT-752 against YFV infection, Syrian golden hamsters were inoculated with a Jimenez hamster-adapted strain, and the prodrug was first administered orally (100, 300 or 1000 mg/kg) 4 h prior to viral challenge, and afterwards as BID doses for 7 consecutive days, beginning 1 h post infection (pi). There was a significant improvement in the survival of the YFV-infected hamsters treated with all doses of AT-752, as compared with the vehicle-treated group (p<0.001; Fig 4 and Table 3). Some mortality was observed in the 1000 and 100 mg/kg treatment groups, while 100% of the 300 mg/kg group survived to the end of the study (Fig 4 and Table 3). Treatment with 1000 mg/kg AT-752 initiated 2 d pi gave comparable survival results to those given the prodrug beginning 4 h prior to the viral challenge, or 50 mg/kg ribavirin (RIBA) which was used as a positive control. A 100% mortality rate (n=15/15) was observed in untreated hamsters infected with YFV, with deaths occurring on Day 6 to 11 pi (Fig 4), resulting in a mean day-to-death of 7.6 ± 1.5. This mortality rate was higher than is typically observed in this model [14].

**Fig 4.**
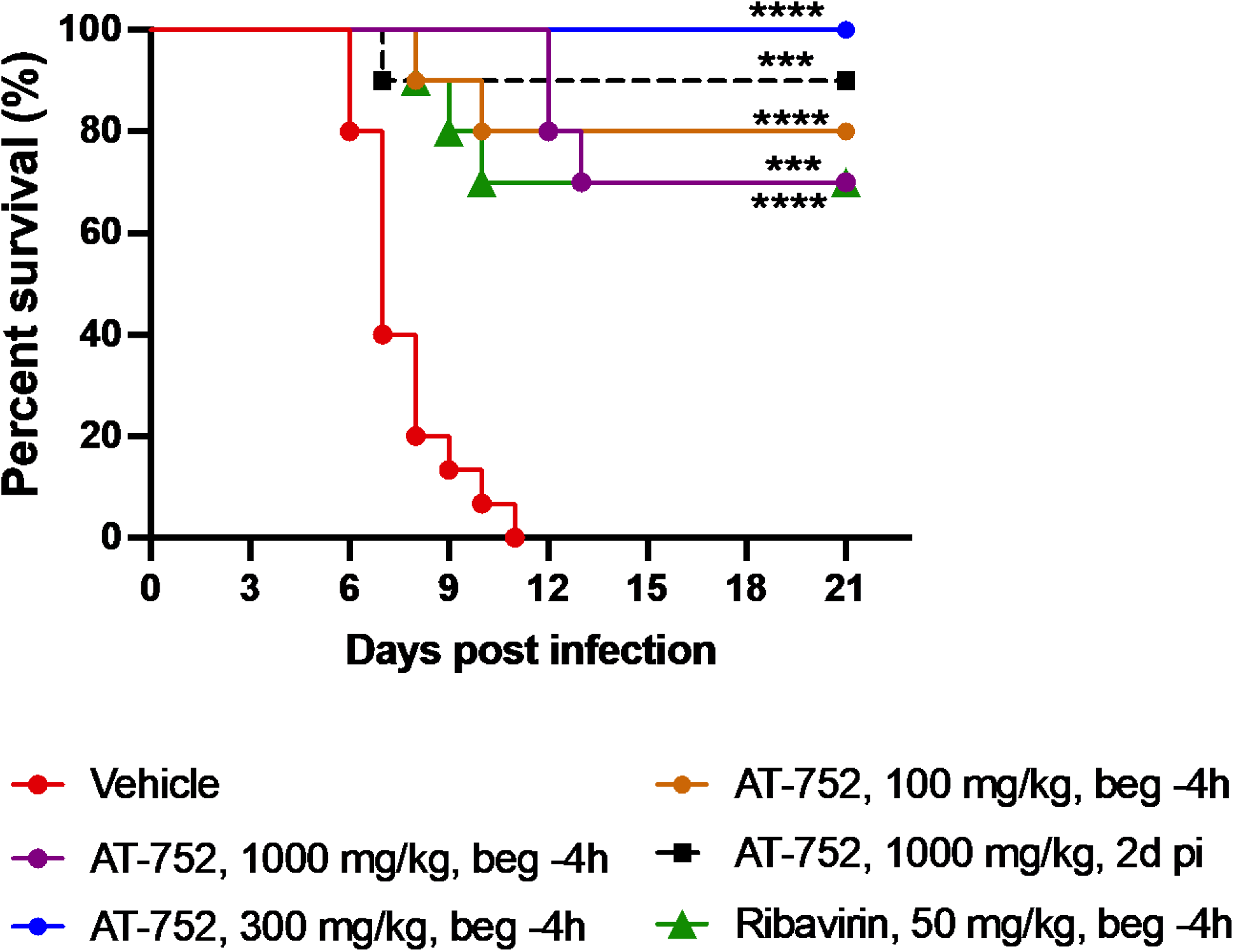
Kaplan-Meier survival curves of YFV-infected hamsters treated with AT-752. Syrian golden hamsters challenged with YFV were administered vehicle (placebo), ribavirin (positive control) or AT-752 four h prior to or 2 d post challenge, followed by BID doses of 1000, 300 or 100 mg/kg for 7 consecutive days starting 1 h post inoculation. Percent survival was calculated up to 21 days post infection. The treated groups were significantly different from vehicle control by one-way ANOVA and Dunnett’s test, ****p<0.0001, ***p<0.001.

**Table 3.**
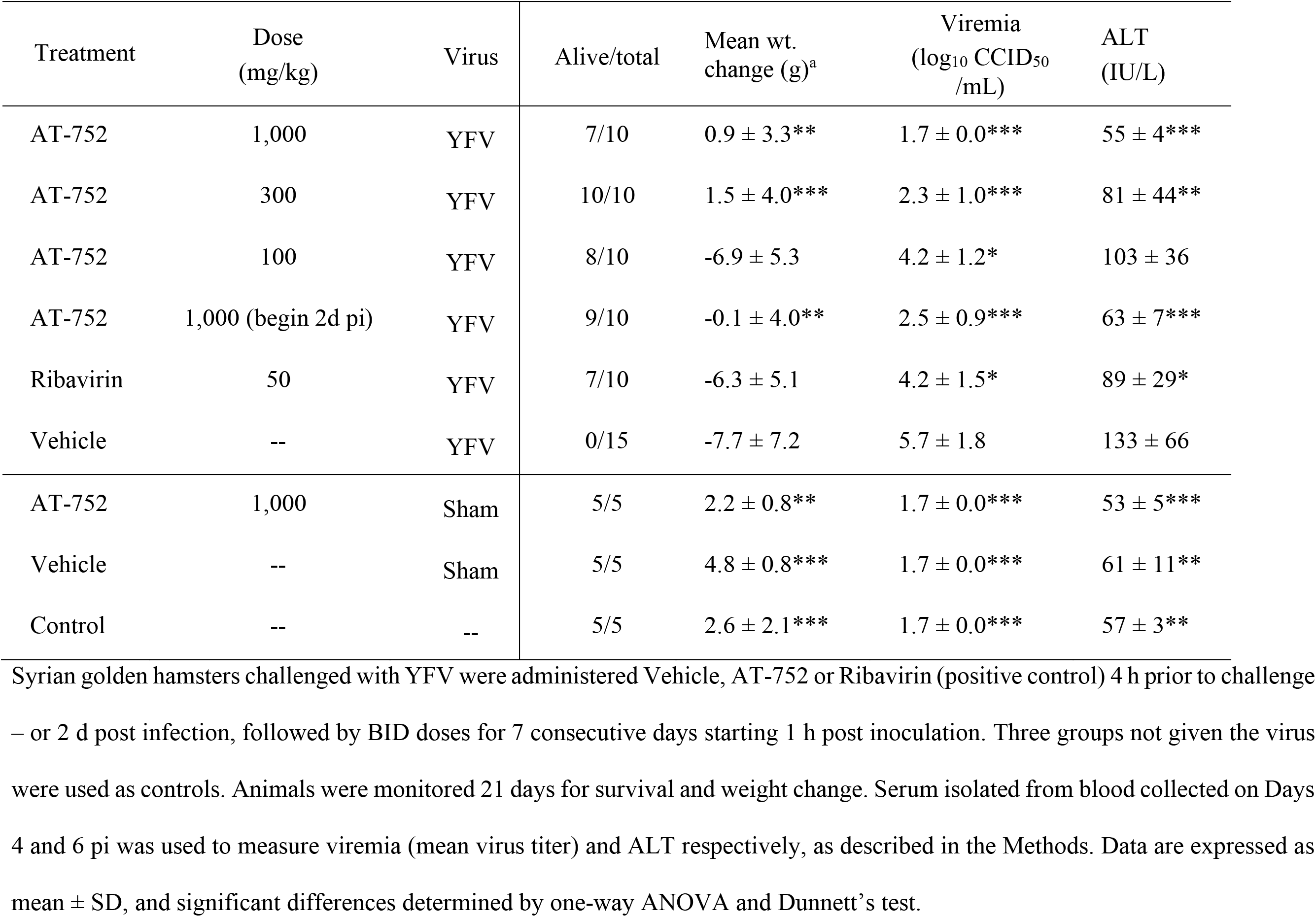
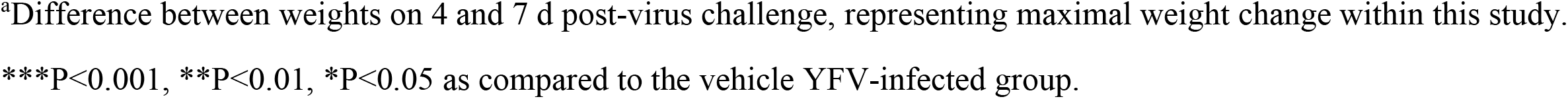
Effects of AT-752 treatment in hamsters infected with yellow fever virus.

| Treatment | Dose<br>(mg/kg) | Virus | Alive/total | Mean wt.<br>change (g)<sup>a</sup> | Viremia<br>(log<sub>10</sub> CCID<sub>50</sub><br>/mL) | ALT<br>(IU/L) |
| --- | --- | --- | --- | --- | --- | --- |
| AT-752 | 1,000 | YFV | 7/10 | 0.9 ± 3.3** | 1.7 ± 0.0*** | 55 ± 4*** |
| AT-752 | 300 | YFV | 10/10 | 1.5 ± 4.0*** | 2.3 ± 1.0*** | 81 ± 44** |
| AT-752 | 100 | YFV | 8/10 | -6.9 ± 5.3 | 4.2 ± 1.2* | 103 ± 36 |
| AT-752 | 1,000 (begin 2d pi) | YFV | 9/10 | -0.1 ± 4.0** | 2.5 ± 0.9*** | 63 ± 7*** |
| Ribavirin | 50 | YFV | 7/10 | -6.3 ± 5.1 | 4.2 ± 1.5* | 89 ± 29* |
| Vehicle | -- | YFV | 0/15 | -7.7 ± 7.2 | 5.7 ± 1.8 | 133 ± 66 |
| AT-752 | 1,000 | Sham | 5/5 | 2.2 ± 0.8** | 1.7 ± 0.0*** | 53 ± 5*** |
| Vehicle | -- | Sham | 5/5 | 4.8 ± 0.8*** | 1.7 ± 0.0*** | 61 ± 11** |
| Control | -- | -- | 5/5 | 2.6 ± 2.1*** | 1.7 ± 0.0*** | 57 ± 3** |

Syrian golden hamsters challenged with YFV were administered Vehicle, AT-752 or Ribavirin (positive control) 4 h prior to challenge – or 2 d post infection, followed by BID doses for 7 consecutive days starting 1 h post inoculation. Three groups not given the virus were used as controls. Animals were monitored 21 days for survival and weight change. Serum isolated from blood collected on Days 4 and 6 pi was used to measure viremia (mean virus titer) and ALT respectively, as described in the Methods. Data are expressed as mean ± SD, and significant differences determined by one-way ANOVA and Dunnett's test.
<sup>a</sup>Difference between weights on 4 and 7 d post-virus challenge, representing maximal weight change within this study.
\*\*\*P<0.001, \*\*P<0.01, \*P<0.05 as compared to the vehicle YFV-infected group.

Body weights of the hamsters were recorded on day 0 and then daily from day 3 to 18 pi to evaluate the effect of treatment on any change as an indicator of health. Weight change curves of the YFV-infected groups treated with 1000 or 300 mg/kg AT-752 were similar to those of sham-infected animals treated with 1000 mg/kg AT-752 (Fig 5). Moreover, there was no significant difference in weight change curves between AT-752-treated animals and normal controls and no toxicity, as evidenced by a lack of weight loss or mortality, was observed from AT-752 treatment in these animals. This is different from the animals treated with 50 mg/kg/d of RIBA (positive control) where weight change declined steeply after Day 6 pi and rebounded after Day 10 pi (Fig 5). There was also a drop in the average weight of infected hamsters treated with 100 mg/kg AT-752 between Days 6 and 9 pi, which diverged from the other AT-752 treatment groups (Fig 5). These weight change curves corresponded with weight change between Days 4 and 7 pi (Table 3), when maximal weight change occurred in this study.

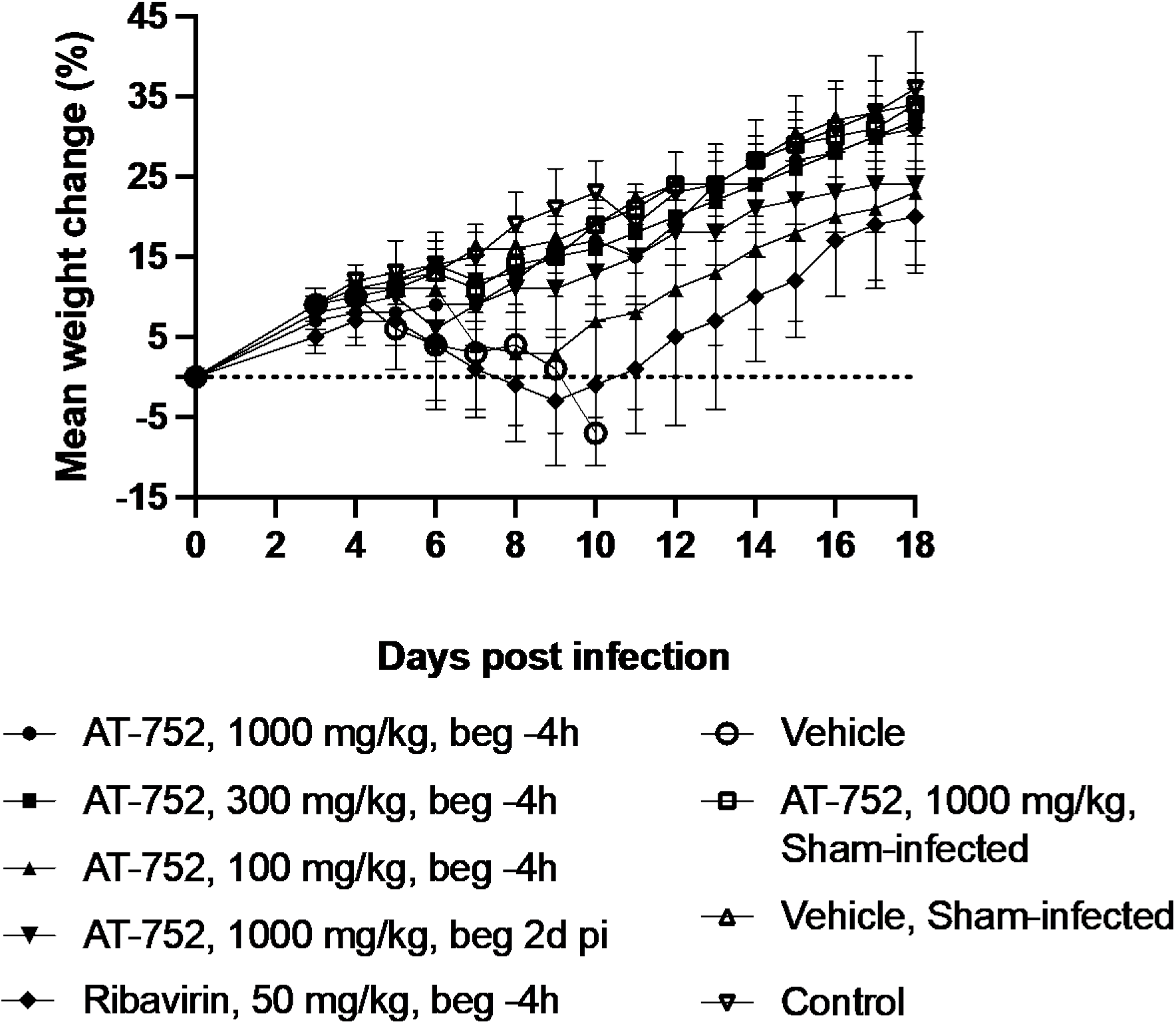

YFV-infected animals treated with 1000 or 300 mg/kg AT-752, regardless of treatment initiation time, had significantly reduced weight loss compared to vehicle control (Table 3; p <0.01 by ANOVA and Dunnett’s test). However, there was no significant improvement in weight change observed in YFV-infected animals treated with 100 mg/kg AT-752 or with 50 mg/kg RIBA (Table 3). Taken together, these data demonstrate a dose-dependent change in weight for groups treated with AT-752. The decline in average weight change of YFV-infected vehicle-treated animals began after Day 4 pi and continued until the animals had succumbed to disease or were humanely euthanized (Fig 5). As expected, sham-infected hamsters, untreated or vehicle-treated, had a consistent increase in weight over the course of the study (Fig 5, Table 3).

Viremia, a primary endpoint, was measured in serum collected on Day 4 pi. For the animals infected with YFV, all the treatment groups had significantly lower viremia titers as compared with vehicle only treatment (Fig 6, Table 2). In fact, the virus titer measured in the 1000 mg/kg AT-752 treated group initiated 4 h prior to virus inoculation was similar to the basal levels measured in the sham-infected control groups – representing the assay’s limit of detection – which was about 4 log_10_-fold lower than the vehicle only group (p<0.001). The drop in viremia occurred even when AT-752 treatment was initiated at 2 dpi, being more than 2 log_10_ lower than the untreated infected animals on Day 4 pi (Fig 6). Higher average titers in animals treated with 100 mg/kg AT-752 or 50 mg/kg RIBA indicated a more modest 1.4 log_10_ decrease in viremia compared to the vehicle only group (p<0.05; Table 2). These data are consistent with survival and weight change data. A similar pattern to the viremia results was observed in ALT concentrations measured from serum collected at Day 6 pi. Except for the group treated with 100 mg/kg AT-752, all the treatment groups had significantly lower ALT levels than the vehicle only group (p<0.05; Table 2). These data showed a dose-dependent improvement in serum ALT when YFV-infected animals were treated with AT-752, regardless of when treatment was initiated.

**Fig 6.**
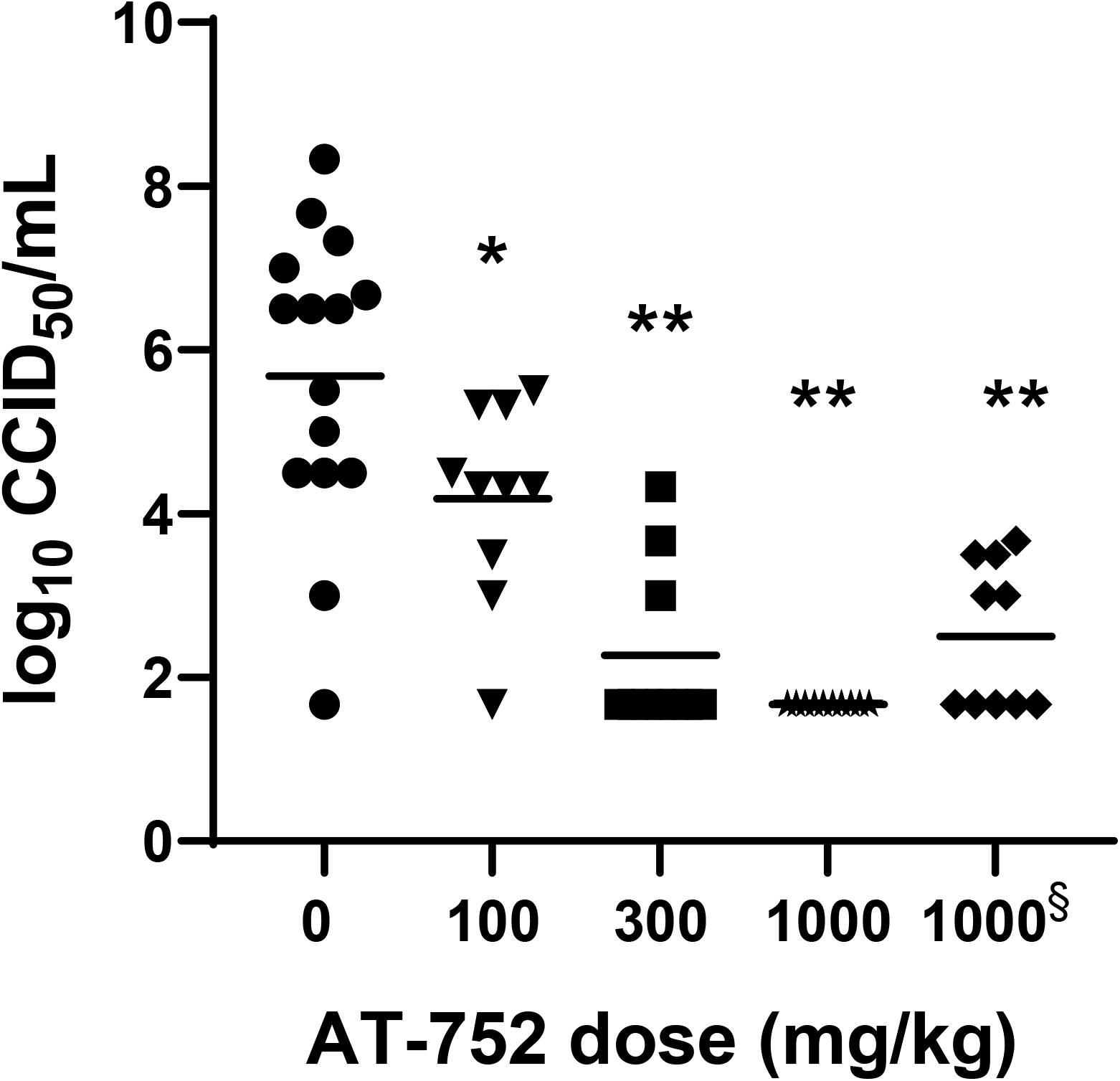
Virus titer in YFV-infected hamsters treated with AT-752. Syrian golden hamsters challenged with YFV were administered 0 (vehicle), 100, 300 or 1000 mg/kg AT-752 four h prior, followed by BID doses for 7 consecutive days starting 1 h post inoculation (pi). Serum was collected 4 d pi, and titers measured by an infectious assay as described in the Methods. The treated groups were significantly different from vehicle by one-way ANOVA and Dunnett’s test, *p<0.05, **p<0.001. § AT-752 dosing began 2 d post challenge.

## Discussion

Yellow fever, which is caused by infection from the mosquito-borne YFV, can lead to a lethal viral hemorrhagic fever associated with liver and kidney failure [2]. With mosquito control and vaccination campaigns, it was no longer considered a dangerous infectious disease by the end of the 20^th^ century but had become a neglected tropical disease. However, in 2016, yellow fever re-emerged as a human threat when outbreaks occurred in endemic areas and in non-endemic areas with historically low YFV activity, possibly aided by climate change and an increase in densely populated areas with low vaccination coverage [5]. Thus, there is an urgent need for a safe and efficacious DAA to aid in the management of these unexpected events that have resulted in significant morbidity and mortality.

Here, we used an animal model to show that oral treatment with AT-752 is effective in improving disease parameters such as survival, serum ALT levels and weight change in hamsters infected with YFV. Pre-treatment with AT-752 (4 h prior to viral challenge) significantly reduced virus load in a dose-dependent manner (Table 2). Just as importantly, when AT-752 treatment was initiated two days after infection, viremia in the infected hamsters was decreased by >2 log_10_ on Day 4 pi, a time noted for clinical symptoms in this model [14]. While this latter result is promising, more work would need to be done to determine the efficacy of AT-752 in treating the disease after clinical symptoms are apparent, a more likely scenario with humans.

YFV belongs to the genus *Flavivirus*, which includes the Dengue and Zika viruses. These viruses have a highly conserved NS5 whose interactions, if disrupted, result in the loss of viral replication, likely across all flaviviruses [4, 15]. We have previously reported that AT-752 has a direct effect on dengue viral replication, and presumably YFV as well, by forming the intracellular active triphosphate AT-9010, which then inhibits the RdRp responsible for viral replication [12]. In addition, the tissue distribution study revealed that the highest concentrations of AT-9010 were found in liver and kidney, the two organs most affected by YFV. These data show that not only does the prodrug significantly reduce viremia in this animal model of disease, but the greatest amounts of the active triphosphate metabolite were formed in the tissues most affected by the virus.

Despite many attempts to develop an effective treatment for yellow fever, including other nucleoside analogs [3, 10, 11, 14], no compound has successfully demonstrated efficacy against the disease in human testing. Sofosbuvir, a clinically approved drug against hepatitis C virus (HCV), has recently been shown to inhibit YFV *in vitro* and *in vivo* [2, 8], and a clinical trial in Brazil to test its efficacy in humans infected with YFV is currently underway [16]. AT-752 has also been shown to be a potent inhibitor of HCV [12], as is its congener AT-527 which forms the same active triphosphate, and was up to 58-fold more potent than sofosbuvir against HCV clinical isolates [13]. To conclude, AT-752, with its broad potent antiviral activity against flaviviruses and favorable safety profile [12], as well as its demonstrated efficacy in an *in vivo* hamster model, is a promising candidate for clinical development for the treatment of yellow fever.

## Acknowledgments

This research was supported in part by contract HHSN272201700041I Task order A11 from the Virology Branch, NIAID, NIH. The authors thank Dr. Kerry-Ann da Costa for her excellent assistance in preparing this manuscript. This study was sponsored by Atea Pharmaceuticals, Inc.

